# Spatial instability affects Key Biodiversity Areas scoping for bumblebees across three decades

**DOI:** 10.1101/2025.03.01.640949

**Authors:** D. Nania, A. Cristiano, M. Mei, D. Michez, M. Di Musciano, R. Cazzolla Gatti, P. Merelli, M. Pacifici, C. Rondinini, P. Cerretti

## Abstract

Identifying potential Key Biodiversity Areas (KBAs) using species occurrence records is crucial for integrating hyperdiverse taxa, such as insects, into the KBA network. To ensure the KBA network supports effective conservation decision-making, established KBAs need to be reassessed every 8 to 12 years. Realistically, reassessments for very high numbers of species will be based also on opportunistic occurrence points. However, how changes in species distribution data availability and completeness in different time periods influence the spatial stability of potential KBAs over time has not been investigated in any taxa yet. We performed potential KBA assessments for 28 species and subspecies of bumblebees in Italy using an expert-validated dataset of distribution data collected between 1990 and 2023, using 10-year cumulative and progressive time series. Our findings reveal that KBA scoping for species with distributional knowledge gaps does not ensure stability and reliability over time. Potential KBA turnover percentages between different time periods can be high (≃133%). While only seven species/subspecies could trigger potential KBAs, their turnover percentage ranged between 28.5% and 50%. We highlight the need for standardized and strategic monitoring of established KBAs and species’ distributions between reassessments. We also discuss strategies to optimize monitoring plans under resource constraints.

## Introduction

Key Biodiversity Areas (KBA) are one of the most recently developed tools to map the distribution of sites hosting unique fractions of biodiversity on our planet (Langhammer et al., 2021). This approach relies on the application of standardized criteria to biodiversity data, allowing rational evaluation of a given site.

Given the increasing consideration of KBAs in conservation policy making worldwide (CBD, 2022; European Commission, 2022; IAEG-SDGs, 2019), several methods have been developed over the last years to accelerate the scoping process of new potential KBAs (Nania et al., 2024a; Spiliopoulou et al., 2024; Farooq et al., 2020). Such methodologies will be crucial to fill the knowledge gap on the distribution of KBAs, which are currently known for a very small fraction of animal and plant taxa (BirdLife International, 2024).

The identification of KBAs requires an estimate of the global population size of the species under examination. The KBA standard accepts different types of data as proxies for the global population size. Area of Habitat (AOH) maps are listed as the second best data type for this purpose (IUCN, 2016). AOH maps show the distribution of the habitat of a species within its known geographic distribution range and elevation limits (Nania et al., 2022; Lumbierres et al., 2022).

Although novel methodologies that apply KBA criteria to large numbers of species will accelerate the KBA mapping process, it remains unclear whether the outcomes of such KBA scoping assessments portray a reliable network of sites whose geographic boundaries will remain stable over time. The Global Standard for the Identification of KBAs (IUCN KBA Standards and Appeals Committee, 2022) states that KBA sites should be reassessed under the criteria that validated the site as KBA at least once every 8–12 years. This ensures that global KBAs always account for changes in biodiversity distribution, increasing the reliability of the KBA network. This issue is particularly relevant for species whose global geographic distribution remains poorly documented. For these groups, new occurrence records from future sampling campaigns are likely to significantly shift the currently known distribution, reshaping it and including or excluding areas due to data deficiencies. In contrast, this “range shift” effect is much less pronounced for species with well-documented distributions, such as birds and mammals (Troudet et al., 2017).

Species diversity of insects is very high, and they play a key role in maintaining ecosystem health. Alarming signs of decline in insect populations have intensified in recent decades (Müller et al., 2024; Dalton et al., 2023; Wagner et al., 2021; Hallmann et al., 2020), affecting ecosystem functioning (Zhou et al., 2023). Thus, the urgency to map KBAs is particularly strong for insects. However, the low understanding of their distribution ranges potentially undermines the reliability of KBA assessments for these species. Furthermore, for insects the issue of retrieving reliable distribution data for global assessments is particularly relevant. Although large databases such as the Global Biodiversity Information Facility (GIBIF) and iNaturalist (https://www.inaturalist.org) may provide high numbers of occurrence points, these databases are not curated and validated by taxonomists and therefore cannot provide robust data for KBA scoping. This is especially true in the case of many insects, including bumblebees, for which many cryptic species cannot be identified based on a single picture (Bossert, 2015; Williams et al., 2012; Williams et al., 2011), as it happens for research grade observations. Current large-scale projects focusing on pollinators, including bumblebees, like the EU Pollinator Monitoring Scheme (European Commission et al., 2024), will have limited spatial coverage, as its goal is to detect global population trends and not spatial trends. Thus, the most reliable source of distribution data for insects currently is represented by curated expert-validated datasets and will remain so for years to come.

We tested the spatial stability of potential KBA’s for 28 species of bumblebees (genus *Bombus*) in Italy from 1990 to 2023 based on an expert-validated dataset of occurrence records. Some species are represented by single subspecies that are geographically isolated and morphologically distinct from other conspecific populations (Nania et al., 2024b). We measured potential KBA extent and turnover based on their habitat distribution and explored changing patterns in the habitat distribution for the examined species. We hypothesized that besides environmental change, KBA scoping can be highly affected by the lack of appropriate monitoring, especially when dealing with elusive and understudied groups. Finally, we discuss possible solutions to promote species monitoring for KBA reassessments over time.

## Methods

### Distribution data

Occurrence data for 28 species of bumblebees occurring in Italy was retrieved from two datasets: the Atlas of European bees (Rasmont et al., 2015), which has been updated with records collected until 2023, and a private dataset with more than 3,000 records of bumblebees in Italy. We extracted all records for which the year of collection was available and only retained records starting from 1990. The resulting dataset, comprising 853,343 occurrence points, was splitted into six subsets of data following two approaches: a progressive moving-window time series and a cumulative time series. In the progressive time series, each subset covers a 10-year span to simulate a reassessment based on the knowledge of the taxa distribution within each time window, following the indications on time intervals between assessments provided by the Global Standard for the Identification of KBAs (IUCN KBA Standards and Appeals Committee, 2022). The first subset includes occurrences recorded from 1990 until 2000. For each of the subsequent subsets, the rolling window shifts by 5 years forward, thus smoothing occurrence data fluctuations between time periods. The last subset covers a 8 year span, including all occurrences in the dataset from 2015 to 2023. In the cumulative time series, each subset of species distribution data includes all the available data prior to the given time period. The first subset includes occurrences recorded from 1990 until 2000. The subsequent subsets cumulatively include data by five-year increment. This allowed us to compare the different outputs produced in KBA analyses based on a cumulative and a progressive time series.

We derived the distribution range limits of the species and subspecies by overlaying their occurrences with a map of administrative areas (https://www.naturalearthdata.com/). To account for potential artifacts caused by the inclusion or exclusion of areas near occurrence points close to administrative borders, we applied a circular buffer with a 5 km radius around each occurrence point before performing the intersection with the administrative areas. An explanatory scheme of how distribution ranges were delineated is available in Supporting information S1.1. Although administrative boundaries are not linked to the real distribution of the species, this is to be considered as an intermediate step. The final distribution data (AOH maps) was refined by mapping the habitat within the range, significantly increasing data accuracy. The habitat within the distribution range was mapped using the Copernicus land cover map. We used the Copernicus land cover (CGLS-LC100) 2019 categories as surrogates for the habitat of the species and linked the species to their habitat following the species-habitat association and elevation limits table from Nania et al. (2024a), where the authors used the same methodology to map potential KBAs for bumblebees in Italy. A table showing how land cover categories were linked to habitats is available in supplementary information (S1.2). The final AOH maps included only the land cover categories that represent suitable habitat for the species within their distribution ranges and elevation limits. Elevation data was retrieved from the Shuttle Radar Topography Mission (USGS EROS Archive, 2019), resampled to the land cover resolution (100m). AOH maps were produced for all species over each time period.

We additionally produced AOH maps for each species using all available records (1990–2023) to validate our habitat models. The validation of the AOH maps was carried out within the Italian borders for the following reasons:

1. The species-habitat associations in Nania et al. (2024b) are based on information on ecological and distributional data specific to Italy (S1.2).
2. Elevation limits may vary slightly across a species’ global range following a latitudinal gradient, which could affect validation results. However, the primary goal of mapping habitats globally was to estimate global population sizes for applying KBA criteria.
3. The KBA scoping was conducted in Italy, making it essential to ensure the validation test’s accuracy in the study area.

### Key Biodiversity Areas scoping

A hypergeometric distribution test was used to validate the AOH maps based on species occurrence points (Dahal et al., 2021). This validation test was conducted only for species with at least 25 available occurrence points, covering 21 of the 28 *Bombus* species. Further details on the AOH map validation are available in supplementary information (S1.3).

The KBA scoping was performed using the non-stationary grid methodology described in Nania et al. (2024a), in World Cylindrical Equal Area projection (ESRI:54034) as required by the Global Standard for the Identification of KBAs (IUCN KBA Standards and Appeals Committee, 2022). This approach avoids missing potential KBAs due to a fixed position of the grid on the map. We implemented a 10×10 km cell grid to scan the geographic surface of Italy to detect potential KBAs. The same cell size was previously used for scoping potential KBAs for insects, reptiles and amphibians (Ascenzi et al., 2025; Nania et al., 2024a; Nania et al., 2024b). We performed the KBA scoping based on four KBA criteria (A1, B1, B2, B3). For criterion A (threatened species), we refer to the IUCN Red List threat status (IUCN, 2024). These criteria can be applied without any additional data other than the AOH maps. We repeated the procedure for all species for all six subsets of data and produced a single potential KBA map for each time period, merging all the KBAs identified under different criteria for each species. We analyzed changes in the potential KBA map across time periods by comparing each period with the previous one. Specifically, we measured the extent of stable potential KBAs (areas already identified as potential KBAs in the previous time period), gained potential KBAs (areas not identified as potential KBAs in the previous time period but present in the examined time period), and lost potential KBAs (areas identified as potential KBAs in the previous time period but absent in the examined time period). Additionally, we calculated the turnover percentage of the potential KBA network, as the sum of the gained and lost areas expressed as a percentage of the total potential KBA area from the previous time period. We then calculated the turnover percentage of the species triggering potential KBAs, and measured changes in the total potential KBA extent triggered by each species over time, as well as changes in the global AOH of the trigger species. The gain and loss of potential Key Biodiversity Areas (KBAs) over time is driven by changes in the Area of Habitat (AOH). These changes can occur within a defined area for two main reasons: the presence/absence of AOH due to existence/lack of occurrence points within a given administrative area during a specific time period, or the expansion/contraction of AOH within the species’ distribution range, which may cause these areas to no longer meet the KBA criteria, as well as to meet new ones. To disentangle these processes for each species, we measured the percentage of potential KBA turnover driven by changes in the presence/absence of AOH within the triggering areas. The remaining percentage represents areas where AOH presence remains consistent over time, but the criteria were not met for the species within the time period due to changes in the AOH across the species’ distribution range.

We assumed the land cover to be constant throughout the whole time period. This assumption was made because fine-resolution land cover data (i.e. 100m resolution) are not available for the full timeframe of the analyses. However, to test for potential macro-level differences in land cover, we performed a sensitivity analysis using the ESA CCI time series spanning 1992 to 2022 (Copernicus Climate Change Service, Climate Data Store 2019). In this analysis, we combined all land cover classes representing only natural vegetation (i.e. classes 50, 60, 61, 62, 70, 71, 72, 80, 81, 82, 90, 100, 110, 120, 121, 122, 130, 140, 150, 151, 152, 153, 160, 170, 180, 200, 201, 202, 210, 220). Then, using the Italy map as a mask, we calculated the percentage difference in natural land cover types between 2022 and 1992.

## Results

We produced eleven sets of AOH maps – six for the progressive and six for the cumulative time series – that are available on the DRYAD repository [ http://datadryad.org/stash/share/qV6r69zrNUPs1RB3LbwFjwRnawO_DuWfk9Sre9_W0yM.]. The number of species in each subset varies, as occurrence records were not available for some species in certain time periods. A presence/absence table indicating which species lacked records for specific time periods is available in Supporting information (S1.5). The hypergeometric test revealed that 17 (80%) of the 21 tested AOH maps performed better than expected under randomness (S1.4).

Species for which the model did not perform better than under randomness are geographically widely distributed and are unlikely to trigger potential KBAs when applying area-based KBA criteria such as A1, B1 and B2 (S1.3).

The sensitivity test using ESA land cover maps showed a very small difference in natural land cover (average −0.02%) between 1992 and 2022 (S1.4). The KBA scoping across progressive time periods showed a dynamic potential KBA network for bumblebees in Italy, with the total extent varying between 1,982 km² and 3,758 km² depending on the time period examined (Figure 1), and an average value of 2,860.4 km². Criteria A1 and B1 were triggered in all six periods, whereas criterion B2 was triggered in four periods, but not during 1995–2005 and 2000–2010. Criterion B3 was never triggered, as there are not enough geographically restricted species in our sample to trigger any potential KBA site in any time period. The turnover percentage of the potential KBA network ranged from 55.4% to 133.7% (Tab 1), with a mean turnover of 71.8%.

**Figure 1.**
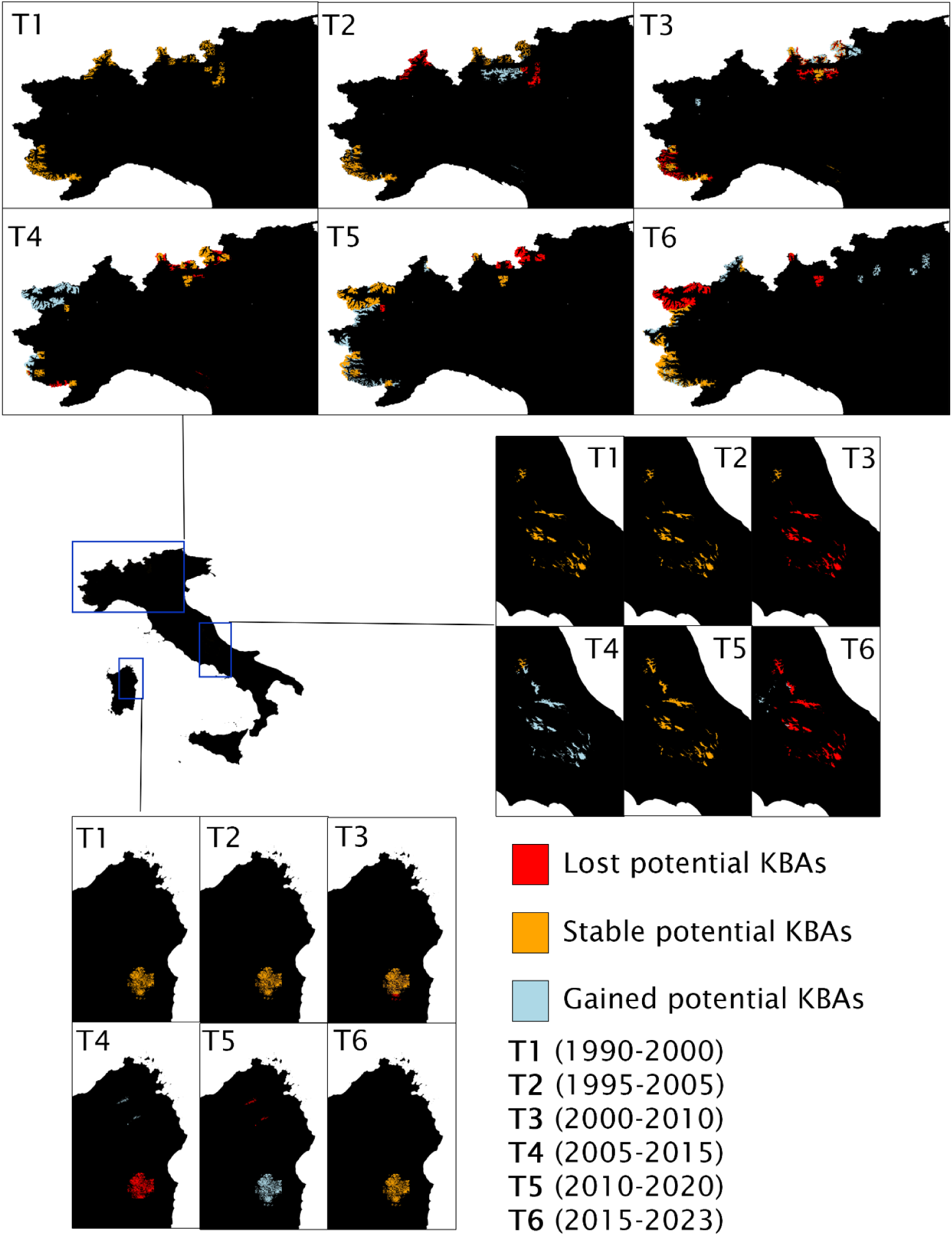
Potential KBAs detected using progressive time series (T1 to T6) regardless of the trigger species and triggered criteria. Yellow areas (Stable KBAs) are potential KBAs that were already detected in the previous time period. Light blue areas (Gained KBAs) are potential KBAs that were detected in the present time period (T1 to T6), but not detected in the previous time period. Red areas (Lost KBAs) are potential KBAs that were detected in the previous time period, but were not detected in the present time period.

For the KBA scoping across cumulative time periods, the total potential KBA extent varies between 2,304 km² and 3,599 km² (Figure 3), with an average value of 2,767.7 km². Criteria A1 and B1 were triggered in all six periods, whereas criterion B2 was triggered only in two time periods (1990-2000 and 1990-2023). The turnover percentage of the potential KBA network ranged from 5.2% to 130.6%, with a mean turnover of 52.8% (Table 1).

**Table 1.** Percentage of potential KBA turnover and species turnover across time periods (T1-T6).

|  | T1 | T2 | T3 | T4 | T5 | T6 |
| --- | --- | --- | --- | --- | --- | --- |
| **Progressive potential KBA turnover** | NA | 55.4% | 91.8% | 133.7% | 66.9% | 80.7% |
| **Cumulative potential KBA turnover** | NA | 28.6% | 78.6% | 5.3% | 21.1% | 130.6% |
| **Progressive species turnover** | NA | 42.9% | 28.6% | 28.6% | 16.7% | 37.5% |
| **Cumulative species turnover** | NA | 33.3% | 28.6% | 0.0% | 0.0% | 37.5% |

Our analysis reveals that gain or loss in KBA cover over time is predominantly driven by changes in AOH within the triggering area. These results are consistent for nearly all species examined. Only two species, *Bombus alpinus helleri* and *Bombus inexspectatus*, deviate from this pattern (S1.6 and S1.7). Seven species triggered potential KBAs across the examined periods (Figure 3). However, not all species triggered potential KBAs in every time period. In the progressive time series, the species turnover between consecutive time periods, measured as the proportion of trigger species either gained or lost relative to the total trigger species across both periods, ranged from 28.5% to 50%, with an average turnover of 38.2% across the six time periods (Table 1). In the cumulative time series, the species turnover was null in two time periods (1990-2015 and 1990-2020), and ranged between 28.5% and 37.5% in the remaining time periods (Table 1).

Potential KBAs extent and AOH extent of single species show a similar trend, as expected (Figure 2). For species with small AOH extents (≤ 5000 km²), positive and negative trends in AOH extent variation across the six time periods are reflected also in the variation of potential KBA extent. For instance, this is the case for *B. alpinus helleri*. However, for species with larger AOH extents (≥ 5000 km²), the AOH extent trend is inverted in the potential KBA extent. This is the case for *B. monticola alpestris* and *B. inexpectatus*.

**Figure 2.**
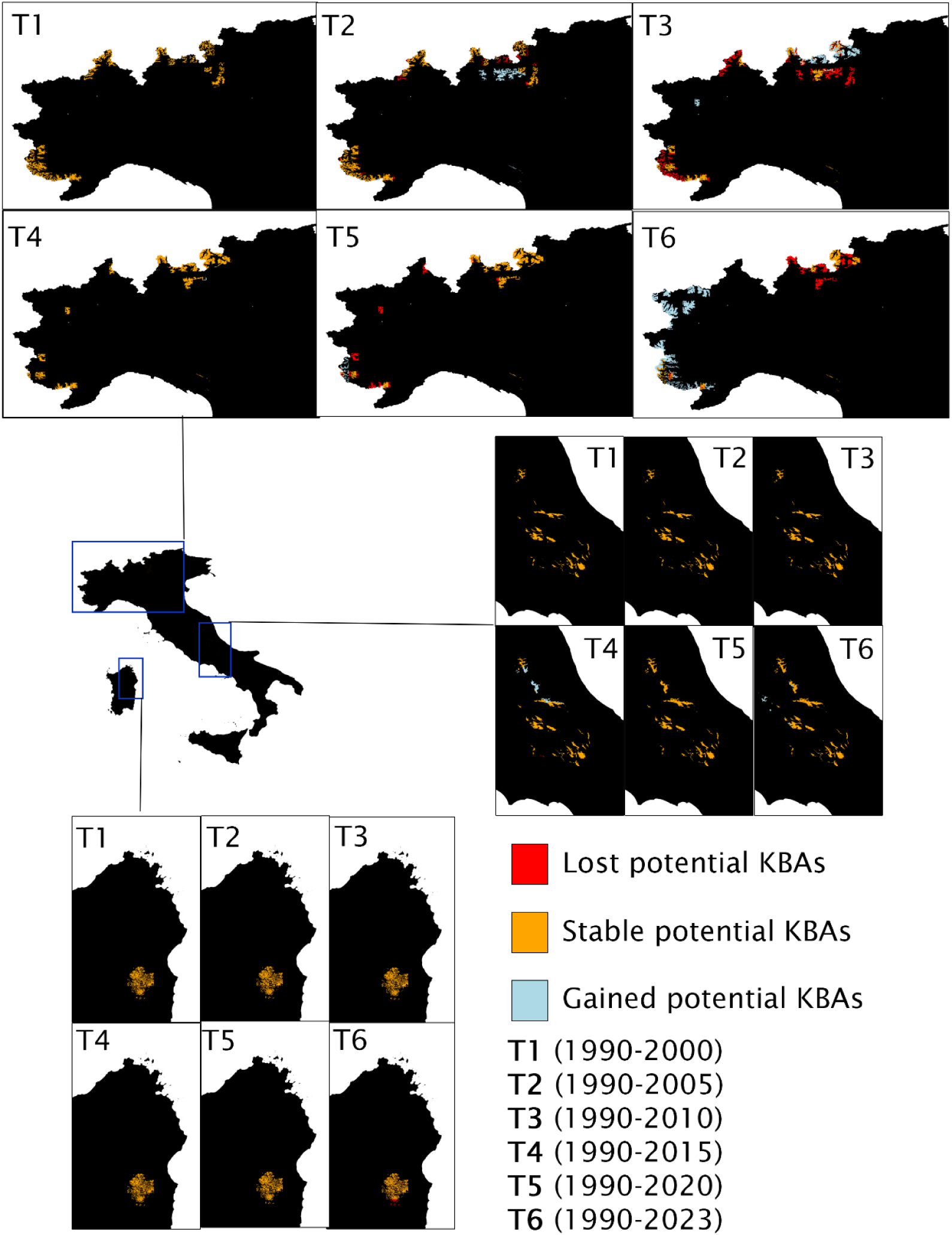
Potential KBAs detected using cumulative time series regardless of the trigger species and triggered criteria. Yellow areas (Stable KBAs) are potential KBAs that were already detected in the previous time period. Light blue areas (Gained KBAs) are potential KBAs that were detected in the present time period, but not detected in the previous time period. Red areas (Lost KBAs) are potential KBAs that were detected in the previous time period, but were not detected in the present time period.

## Discussion

The reliability of the KBA network over time is crucial for its use in supporting spatial planning and conservation strategies (IUCN KBA Standards and Appeals Committee of the IUCN SSC/WCPA). As the number of assessed species is growing (Becker et al., 2024; Spiliopoulou et al., 2024; Farooq et al., 2020), established KBA reassessment will be based also on species occurrence datasets similar to those used in KBA scoping. Re-assessments will be performed by the national coordination groups (IUCN KBA Standards and Appeals Committee of the IUCN SSC/WCPA), which are currently not present in all countries where species occur and will inevitably rely on the species knowledge at the time of the re-assessment.

In this study, the observed fluctuations of the total extent of potential KBAs in both progressive and cumulative time series highlight significant instability of the network over time. In some cases, the percentage of total potential KBA area turnover is extremely high, reaching up to 133.7%. The KBA scoping aligns with previously observed trends (Nania et al., 2024a), confirming that geographically restricted species are more likely to trigger potential KBAs when assessed using specific criteria. This trend was consistent across all assessed time periods. Although only seven species trigger the criteria (Figure 3), the turnover percentage of trigger species and subspecies between time periods can reach 50% (Table 1). As expected, the species-specific contractions and expansions in the extent of habitat are directly mirrored in the extent of potential KBA areas (Figure 3). Potential KBA sites subjected to turnover were shown to be caused by sudden absence or presence of AOH between time periods (S1.6). For all species and subspecies in both progressive and cumulative time series, potential KBA extent increases with the AOH extent up to a threshold, which varies depending on the global AOH extent and distribution. Beyond this threshold, potential KBA extent drops to zero, because the species has become more common and unable to trigger potential KBAs. This was the case for *B. inexpectatus* in both time series.

**Figure 3.**
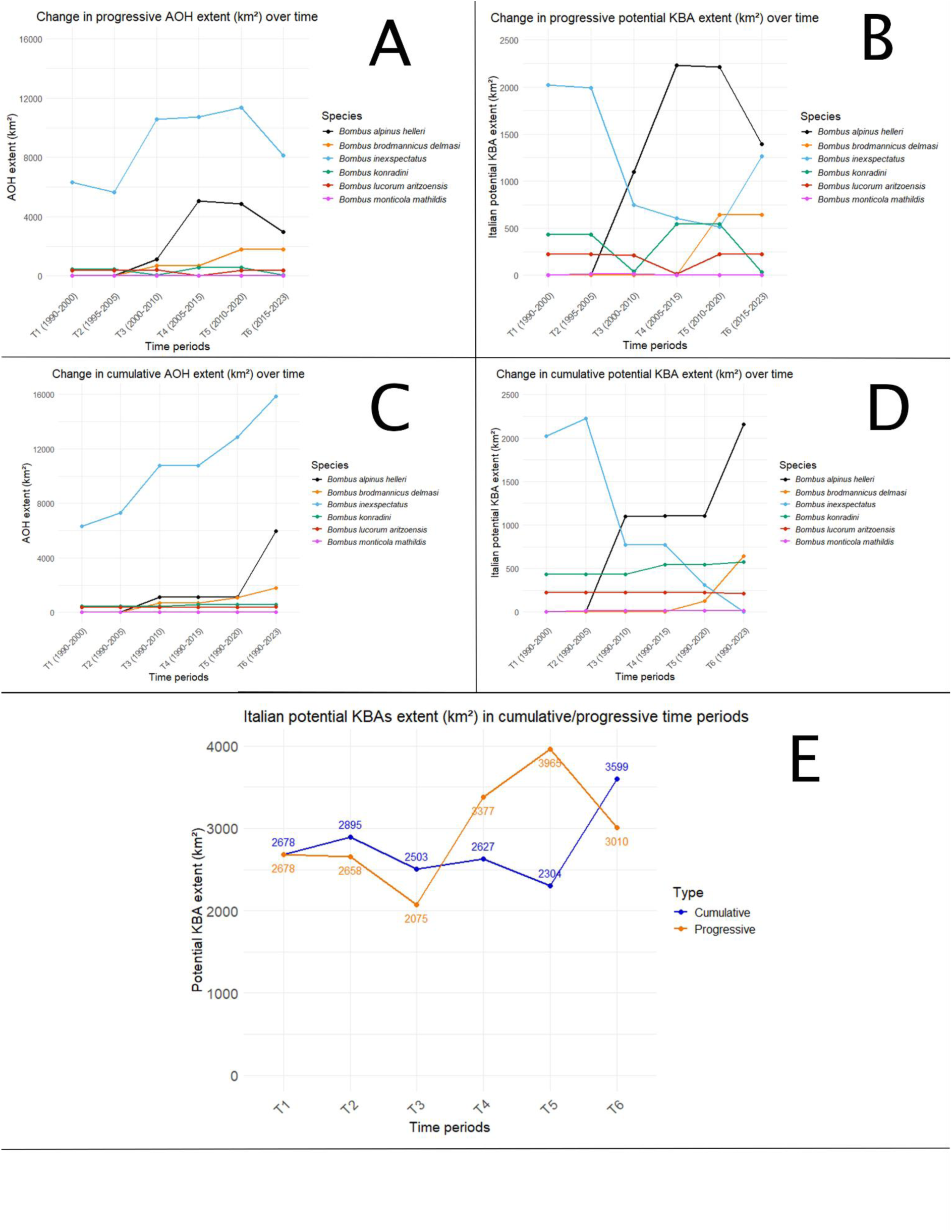
Changes in A) progressive AOH extent; B) progressive potential KBAs extent of single species C) cumulative AOH extent; D) cumulative potential KBAs extent of single species; E) total potential KBA extent for progressive and cumulative time series.

This pronounced instability in the potential KBA network is likely driven by the inherently sporadic nature of occurrence data used to reconstruct species’ distribution ranges (Nania et al., 2024a; Nania et al., 2024b; Spiliopoulou et al., 2024; Farooq et al., 2020). Occurrence points datasets across the species’ distribution range often show temporal and spatial bias (Bowler et al., 2022; Boakes et al., 2010). This issue may be less pronounced for certain vertebrate groups, such as birds, for which global geographic distribution data are typically more accurate and frequent (Troudet et al., 2017), but it is particularly relevant for elusive species like most insects (Isaac et al., 2015; Diniz-Filho et al., 2010). Certain areas tend to be explored more intensively than others during specific historical periods due to various factors, including the provenance of data—whether from museum collections, private collections, or citizen naturalists’ observations. In this context, high variability in the location and extent of KBAs over time is more plausibly a consequence of sampling bias than of actual changes in species’ geographical ranges. Our results highlight the urgent need to establish appropriate and consistent monitoring plans to minimize the risk of mapping potential KBAs that are continually reshaped by data gaps rather than ecological realities.

Although both approaches (progressive and cumulative) adopted here to assess changes in potential KBAs distribution showed high turnover values (Table 1), the cumulative time series overall showed lower turnover percentages compared to the progressive time series, aside from the last time period (T6). Thus, fluctuations in the total potential KBA network are mitigated by the inclusion of older records in the assessment. For conservation planning applications, the cumulative approach may provide a more realistic picture of the actual distribution of the species. Most insect species are likely not uniformly monitored across their whole distribution range (Sánchez-Fernández et al., 2022; Lobo et al., 2007). Low intensity sampling efforts of insect species tend to underestimate species richness (Burner et al., 2021). Therefore, assuming that a species is not present in some areas using solely occurrence points collected within a 10-year time window can lead to weak assessments if not supported by appropriate monitoring plans.

Conducting field sampling to achieve reliable reassessments may involve significant costs (Morant et al., 2020), which are likely to rise as more KBAs are identified and integrated into the global KBA network. In a scenario with limited resources for reassessments and monitoring of species populations, it will be crucial to maximize the efficiency of reassessment by focusing field monitoring efforts on areas most meaningful to the species’ populations. In our three-decade assessment, we highlighted in red (Figure 1 and 2) all potential KBAs that were lost in each time period but were present in the previous one. These areas can play a critical role in the reassessment process by helping prioritize sampling effort in regions where occurrence points are currently missing but where KBAs were identified in the past.

## Conclusion

The ability to identify potential KBAs for hyperdiverse taxa in the coming years— through the implementation of recently developed scoping tools (Nania et al., 2024a; Spiliopoulou et al., 2024; Farooq et al., 2020)— presents an opportunity to use these assessments as a complementary tool to support the reassessment of established KBAs after 8-12 years. This is especially relevant when the number of KBA trigger species is high, and resources for global population monitoring are limited.

Developing strategic monitoring plans will be crucial to collect species distribution data that can underpin reliable reassessments. While field sampling remains indispensable for reassessing established KBAs, KBA scoping can provide valuable insights to optimize these efforts. For example they can highlight areas where sampling is most needed, as demonstrated in this study. In light of these considerations, we believe that future studies should focus not only on including a greater number of species in KBA assessments but also on ensuring the practical maintenance of the KBA network through efficient and well-informed reassessments.

## Acknowledgements

This work received support from The European Union–NextGenerationEU as part of the National Biodiversity Future Center, Italian National Recovery and Resilience Plan (NRRP) Mission 4 Component 2 Investment 1.4 (CUP: B83C22002950007); by the Italian Ministry of University and Research (MUR) under the PRIN Project “Identification of priority sites for the conservation of terrestrial animal and plant diversity to meet European and CBD 2030 targets” CUP B53D23011990006.

