## Supplementary material for "Spatial instability affects Key Biodiversity Areas scoping for bumblebees across three decades": SX Bombus2 - Copia.pdf

**S1.1**. Explanatory scheme of how the range maps were obtained.

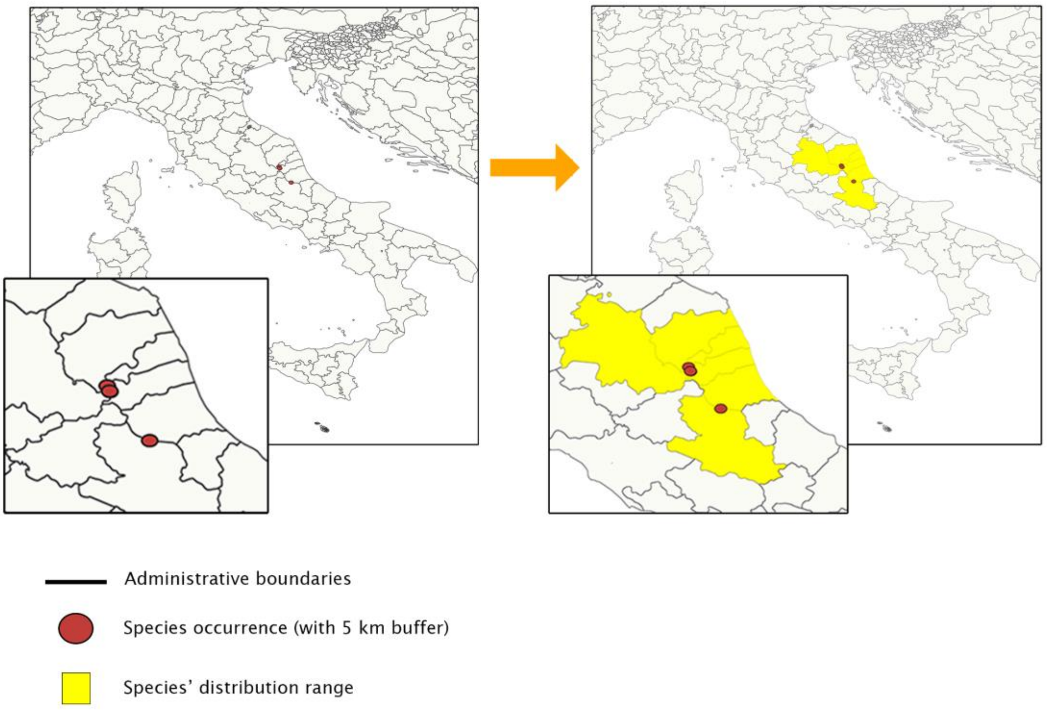

**S1.2** Land cover categories selected as habitat surrogates for each species. A score equal to 0 corresponds to absence of habitat for the species, a score equal to 1 indicates presence of habitat for the species.

| Species | 111 | 115 | 114 | 124 | 20 | 30 | 90 | 50 | 70 | 80 | 60 | 40 | 121 | 126 | 116 | min | max |
|---|---|---|---|---|---|---|---|---|---|---|---|---|---|---|---|---|---|
| *Bombus alpinus helleri* | 0 | 0 | 0 | 0 | 0 | 1 | 0 | 0 | 1 | 0 | 1 | 0 | 0 | 0 | 0 | 2200 | 3100 |
| *Bombus argillaceus* | 0 | 1 | 0 | 1 | 1 | 1 | 0 | 0 | 0 | 0 | 0 | 1 | 1 | 1 | 0 | 0 | 1400 |
| *Bombus barbutellus* | 0 | 0 | 0 | 1 | 1 | 1 | 0 | 0 | 0 | 0 | 0 | 0 | 1 | 1 | 0 | 0 | 2100 |
| *Bombus bohemicus* | 1 | 0 | 1 | 0 | 0 | 0 | 0 | 0 | 0 | 0 | 0 | 0 | 0 | 0 | 1 | 550 | 2300 |
| *Bombus brodmannicus delmasi* | 0 | 0 | 0 | 0 | 0 | 1 | 0 | 0 | 0 | 0 | 0 | 0 | 0 | 0 | 0 | 1500 | 2200 |
| *Bombus campestris* | 0 | 0 | 0 | 1 | 0 | 1 | 0 | 0 | 0 | 0 | 0 | 0 | 1 | 0 | 0 | 200 | 2600 |
| *Bombus cryptarum* | 0 | 1 | 0 | 1 | 0 | 1 | 1 | 0 | 0 | 0 | 0 | 0 | 1 | 0 | 0 | 850 | 2600 |
| *Bombus flavidus* | 0 | 0 | 0 | 1 | 0 | 1 | 1 | 0 | 0 | 0 | 1 | 0 | 1 | 0 | 0 | 1600 | 2700 |
| *Bombus gerstaeckeri* | 0 | 0 | 0 | 0 | 0 | 1 | 1 | 0 | 0 | 0 | 0 | 0 | 0 | 0 | 0 | 900 | 2100 |
| *Bombus inexspectatus* | 0 | 0 | 0 | 0 | 0 | 1 | 0 | 0 | 0 | 0 | 1 | 0 | 0 | 0 | 0 | 1100 | 2400 |
| *Bombus konradini* | 0 | 0 | 0 | 0 | 0 | 1 | 0 | 0 | 0 | 0 | 1 | 0 | 0 | 0 | 0 | 1700 | 2350 |
| *Bombus lapidarius* | 0 | 0 | 0 | 1 | 1 | 1 | 0 | 0 | 0 | 0 | 0 | 1 | 1 | 0 | 0 | 50 | 2850 |
| *Bombus lucorum aritzoensis* | 0 | 0 | 0 | 0 | 0 | 1 | 0 | 0 | 0 | 0 | 0 | 0 | 0 | 1 | 0 | 900 | 1750 |
| *Bombus mendax* | 0 | 0 | 0 | 0 | 0 | 1 | 0 | 0 | 0 | 0 | 0 | 0 | 1 | 0 | 0 | 1300 | 2900 |
| *Bombus mesomelas* | 0 | 0 | 0 | 0 | 0 | 1 | 0 | 0 | 0 | 0 | 1 | 0 | 0 | 0 | 0 | 1000 | 2700 |
| *Bombus monticola alpestris* | 0 | 0 | 0 | 1 | 0 | 1 | 0 | 0 | 0 | 0 | 0 | 0 | 1 | 0 | 0 | 800 | 3100 |

| | | | | | | | | | | | | | | | | | |
|---|---|---|---|---|---|---|---|---|---|---|---|---|---|---|---|---|---|
| *Bombus monticola mathildis* | 0 | 0 | 0 | 1 | 0 | 1 | 0 | 0 | 0 | 0 | 0 | 0 | 1 | 0 | 0 | 1500 | 1800 |
| *Bombus mucidus* | 0 | 0 | 0 | 0 | 0 | 1 | 0 | 0 | 0 | 0 | 1 | 0 | 0 | 0 | 0 | 1200 | 2850 |
| *Bombus norvegicus* | 1 | 0 | 1 | 1 | 0 | 0 | 0 | 0 | 0 | 0 | 0 | 0 | 1 | 0 | 0 | 1000 | 2300 |
| *Bombus pratorum* | 1 | 1 | 1 | 1 | 0 | 0 | 0 | 0 | 0 | 0 | 0 | 0 | 1 | 0 | 0 | 0 | 2600 |
| *Bombus pyrenaeus* | 0 | 0 | 0 | 0 | 0 | 1 | 0 | 0 | 0 | 0 | 1 | 0 | 1 | 0 | 0 | 1400 | 2700 |
| *Bombus quadricolor* | 0 | 0 | 0 | 1 | 0 | 0 | 0 | 0 | 0 | 0 | 0 | 0 | 1 | 0 | 0 | 1000 | 1800 |
| *Bombus ruderatus* | 0 | 0 | 0 | 1 | 1 | 1 | 1 | 0 | 0 | 0 | 0 | 1 | 0 | 1 | 0 | 0 | 2100 |
| *Bombus rupestris* | 0 | 0 | 0 | 1 | 1 | 1 | 0 | 0 | 0 | 0 | 0 | 1 | 1 | 0 | 0 | 300 | 2800 |
| *Bombus subterraneus* | 0 | 0 | 0 | 0 | 1 | 1 | 0 | 0 | 0 | 0 | 0 | 0 | 0 | 0 | 0 | 500 | 2100 |
| *Bombus sylvestris* | 1 | 1 | 1 | 1 | 0 | 0 | 0 | 0 | 0 | 0 | 0 | 0 | 1 | 0 | 0 | 300 | 2500 |
| *Bombus vestalis* | 0 | 1 | 0 | 1 | 1 | 0 | 0 | 0 | 0 | 0 | 0 | 0 | 0 | 1 | 0 | 0 | 2100 |
| *Bombus wurflenii* | 1 | 1 | 0 | 1 | 0 | 0 | 0 | 0 | 0 | 0 | 0 | 0 | 1 | 0 | 0 | 300 | 2700 |

A detailed description of the Copernicus land cover classes is available in the user manual on the official land cover website:

https://land.copernicus.eu/en/products/global-dynamic-land-cover/copernicus-global-land-service-land-cover-100m-collection-3-epoch-2019-globe#download

**S1.3**. Details of the hypergeometric distribution test

| Species name | Confidence interval | Number of points | Upper limit | Lower limit | Number of points inside the model | Upper limit percentage | Point prevalence | Model prevalence |
|---|---|---|---|---|---|---|---|---|
| *Bombus alpinus helleri* | 1.35 | 4 | 2 | 0 | 3 | 50% | 75% | 19.23% |
| *Bombus argillaceus* | 1.35 | 132 | 78 | 56 | 86 | 59.09% | 65.15% | 50.67% |
| *Bombus barbutellus* | 1.35 | 115 | 48 | 29 | 56 | 41.74% | 48.70% | 33.35% |
| *Bombus bohemicus* | 1.35 | 88 | 35 | 18 | 37 | 39.77% | 42.05% | 29.67% |
| *Bombus brodmannicus delmasi* | 1.35 | 1 | 1 | 0 | 0 | 100% | 0% | 9.22% |
| *Bombus campestris* | 1.35 | 89 | 18 | 6 | 19 | 20.22% | 21.35% | 13.08% |
| *Bombus cryptarum* | 1.35 | 31 | 11 | 3 | 20 | 35.48% | 64.52% | 21.84% |
| *Bombus flavidus* | 1.35 | 12 | 6 | 0 | 8 | 50% | 66.67% | 22.61% |
| *Bombus gerstaeckeri* | 1.35 | 37 | 7 | 0 | 12 | 18.92% | 32.43% | 8.69% |
| *Bombus inexspectatus* | 1.35 | 31 | 9 | 1 | 21 | 29.03% | 67.74% | 15.23% |
| *Bombus konradini* | 1.35 | 12 | 2 | 0 | 10 | 16.67% | 83.33% | 2.77% |
| *Bombus lapidarius* | 1.35 | 361 | 146 | 110 | 121 | 40.44% | 33.52% | 35.44% |
| *Bombus lucorum aritzoensis* | 1.35 | 6 | 2 | 0 | 1 | 33.33% | 16.67% | 4.33% |
| *Bombus mendax* | 1.35 | 71 | 24 | 10 | 57 | 33.80% | 80.28% | 24.11% |
| *Bombus mesomelas* | 1.35 | 158 | 31 | 14 | 81 | 19.62% | 51.27% | 13.91% |
| *Bombus monticola alpestris* | 1.35 | 48 | 17 | 6 | 28 | 35.42% | 58.33% | 23.93% |
| *Bombus monticola mathildis* | 1.35 | 1 | 0 | 0 | 0 | 0% | 0% | 0.26% |
| *Bombus mucidus* | 1.35 | 76 | 20 | 7 | 54 | 26.32% | 71.05% | 17.74% |
| *Bombus norvegicus* | 1.35 | 15 | 6 | 0 | 2 | 40% | 13.33% | 19.44% |
| *Bombus pratorum* | 1.35 | 345 | 132 | 98 | 109 | 38.26% | 31.59% | 33.38% |
| *Bombus pyrenaeus* | 1.35 | 96 | 28 | 12 | 65 | 29.17% | 67.71% | 20.68% |
| *Bombus quadricolor* | 1.35 | 31 | 1 | 0 | 0 | 3.23% | 0% | 0.69% |
| *Bombus ruderatus* | 1.35 | 116 | 85 | 65 | 58 | 73.28% | 50% | 64.86% |
| *Bombus rupestris* | 1.35 | 120 | 35 | 17 | 48 | 29.17% | 40% | 21.56% |
| *Bombus subterraneus* | 1.35 | 59 | 9 | 1 | 10 | 15.25% | 16.95% | 7.92% |

| | | | | | | | | |
|---|---|---|---|---|---|---|---|---|
| *Bombus sylvestris* | 1.35 | 109 | 45 | 26 | 47 | 41.28% | 43.12% | 32.13% |
| *Bombus vestalis* | 1.35 | 82 | 26 | 11 | 27 | 31.71% | 32.93% | 22.69% |
| *Bombus wurflenii* | 1.35 | 114 | 26 | 10 | 30 | 22.81% | 26.32% | 15.52% |

**S1.4**. Sensitivity analysis on natural land cover classes using the ESA CCI time series spanning 1992 to 2022.

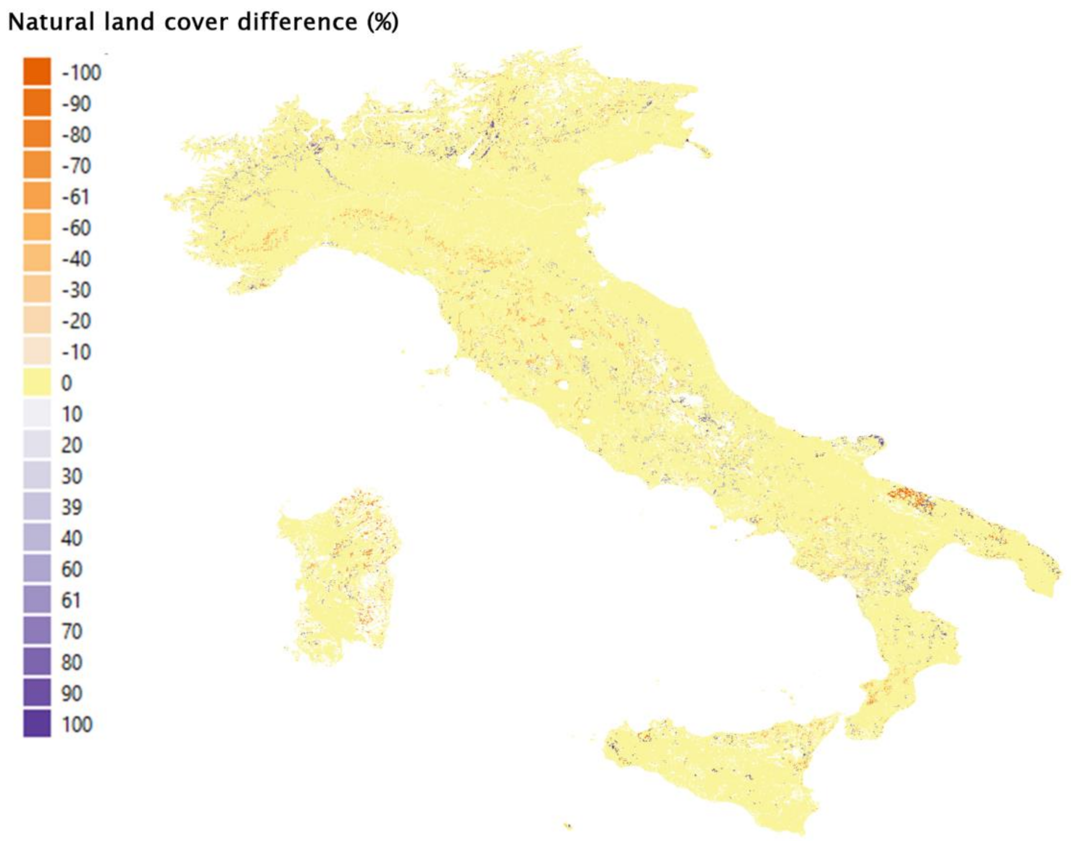

**S1.5**. Binary presence/absence table showing which species lacked occurrence records within the time periods.

| Species | T1 | T2 | T3 | T4 | T5 | T6 |
|---|---|---|---|---|---|---|
| *Bombus alpinus helleri* | 0 | 0 | 1 | 1 | 1 | 1 |
| *Bombus argillaceus* | 1 | 1 | 1 | 1 | 1 | 1 |
| *Bombus barbutellus* | 1 | 1 | 1 | 1 | 1 | 1 |
| *Bombus bohemicus* | 1 | 1 | 1 | 1 | 1 | 1 |
| *Bombus brodmannicus delmasi* | 0 | 0 | 1 | 1 | 1 | 1 |
| *Bombus campestris* | 1 | 1 | 1 | 1 | 1 | 1 |
| *Bombus cryptarum* | 1 | 1 | 1 | 1 | 1 | 1 |
| *Bombus flavidus* | 1 | 1 | 1 | 1 | 1 | 1 |
| *Bombus gerstaeckeri* | 1 | 1 | 1 | 1 | 1 | 1 |
| *Bombus inexspectatus* | 1 | 1 | 1 | 1 | 1 | 1 |
| *Bombus konradini* | 1 | 1 | 1 | 1 | 1 | 1 |
| *Bombus lapidarius* | 1 | 1 | 1 | 1 | 1 | 1 |
| *Bombus lucorum aritzoensis* | 1 | 1 | 1 | 1 | 1 | 1 |
| *Bombus mendax* | 1 | 1 | 1 | 1 | 1 | 1 |
| *Bombus mesomelas* | 1 | 1 | 1 | 1 | 1 | 1 |
| *Bombus monticola alpestris* | 1 | 1 | 1 | 1 | 1 | 1 |
| *Bombus monticola mathildis* | 0 | 1 | 1 | 0 | 0 | 0 |
| *Bombus mucidus* | 1 | 1 | 1 | 1 | 1 | 1 |
| *Bombus norvegicus* | 1 | 1 | 1 | 1 | 1 | 1 |
| *Bombus pratorum* | 1 | 1 | 1 | 1 | 1 | 1 |
| *Bombus pyrenaeus* | 1 | 1 | 1 | 1 | 1 | 1 |
| *Bombus quadricolor* | 1 | 1 | 1 | 1 | 1 | 1 |
| *Bombus ruderatus* | 1 | 1 | 1 | 1 | 1 | 1 |
| *Bombus rupestris* | 1 | 1 | 1 | 1 | 1 | 1 |
| *Bombus subterraneus* | 1 | 1 | 1 | 1 | 1 | 1 |

| | | | | | | |
|---|---|---|---|---|---|---|
| *Bombus sylvestris* | 1 | 1 | 1 | 1 | 1 | 1 |
| *Bombus vestalis* | 1 | 1 | 1 | 1 | 1 | 1 |
| *Bombus wurflenii* | 1 | 1 | 1 | 1 | 1 | 1 |

**S1.6.** Percentage of gained/lost AOH within areas triggering potential KBAs per species and time period in the progressive time series.

| Total potential KBA ( km²) | Gained potential KBA ( km²) | Lost potential KBA ( km²) | % of gained AOH | % of lost AOH | name | Time period |
|---|---|---|---|---|---|---|
| 2709.9 | 687.4 | 718.9 | 0.8 | 0.7 | *Bombus_inexspectatus* | T1 |
| 430.5 | 0 | 0 | 0 | 0 | *Bombus_konradini* | T1 |
| 238.8 | 13.4 | 13.8 | 1 | 0 | *Bombus_lucorum_aritzoensis* | T1 |
| 129.4 | 0 | 129.4 | 0 | 0 | *Bombus_monticola_alpestris* | T1 |
| 11.6 | 11.6 | 0 | 1 | 0 | *Bombus_monticola_mathildis* | T1 |
| 2092.4 | 101.4 | 1342.3 | 0.9 | 0 | *Bombus_inexspectatus* | T2 |
| 432.7 | 2.2 | 395 | 1 | 1 | *Bombus_konradini* | T2 |
| 255.4 | 30.4 | 43.2 | 1 | 0 | *Bombus_lucorum_aritzoensis* | T2 |
| 11.6 | 0 | 0 | 0 | 0 | *Bombus_monticola_mathildis* | T2 |
| 1095.3 | 1095.3 | 0 | 1 | 0 | *Bombus_alpinus_helleri* | T2 |
| 0.1 | 0.1 | 0 | 1 | 0 | *Bombus_brodmannicus_delmasi* | T2 |
| 2669.8 | 1574.4 | 436.4 | 0.9 | 0 | *Bombus_alpinus_helleri* | T3 |
| 0.1 | 0 | 0 | 0 | 0 | *Bombus_brodmannicus_delmasi* | T3 |
| 750.1 | 0.1 | 141.9 | 1 | 0 | *Bombus_inexspectatus* | T3 |
| 547.6 | 509.8 | 2.1 | 1 | 0 | *Bombus_konradini* | T3 |
| 226.1 | 13.9 | 212.3 | 0.4 | 1 | *Bombus_lucorum_aritzoensis* | T3 |
| 11.6 | 0 | 11.6 | 0 | 1 | *Bombus_monticola_mathildis* | T3 |
| 2920.8 | 687.3 | 710.1 | 0.9 | 0.8 | *Bombus_alpinus_helleri* | T4 |
| 644.8 | 644.7 | 0.1 | 1 | 0 | *Bombus_brodmannicus_delmasi* | T4 |
| 678.6 | 70.4 | 165.7 | 0.9 | 0.3 | *Bombus_inexspectatus* | T4 |

| | | | | | | |
|---|---|---|---|---|---|---|
| 553.9 | 8.4 | 7.6 | 0.9 | 0 | *Bombus_konradini* | T4 |
| 239.3 | 225.4 | 13.9 | 0 | 1 | *Bombus_lucorum_aritzoensis* | T4 |
| 2647.5 | 436.8 | 1253.8 | 0.2 | 0.9 | *Bombus_alpinus_helleri* | T5 |
| 644.7 | 0 | 0 | 0 | 0 | *Bombus_brodmannicus_delmasi* | T5 |
| 1501.5 | 988.6 | 235.2 | 0.2 | 0.8 | *Bombus_inexspectatus* | T5 |
| 581.6 | 35.3 | 545.9 | 1 | 1 | *Bombus_konradini* | T5 |
| 235 | 9.5 | 9.7 | 1 | 0 | *Bombus_lucorum_aritzoensis* | T5 |
| 1393.6 | 0 | 1393.6 | 0 | 1 | *Bombus_alpinus_helleri* | T6 |
| 644.7 | 0 | 644.7 | 0 | 1 | *Bombus_brodmannicus_delmasi* | T6 |
| 1266.3 | 0 | 1266.3 | 0 | 1 | *Bombus_inexspectatus* | T6 |
| 35.7 | 0 | 35.7 | 0 | 1 | *Bombus_konradini* | T6 |
| 225.3 | 0 | 225.3 | 0 | 1 | *Bombus_lucorum_aritzoensis* | T6 |

**S1.7.** Percentage of gained/lost AOH within areas triggering potential KBAs per species and time period in the cumulative time series.

| Total potential KBA ( km²) | Gained potential KBA ( km²) | Lost potential KBA ( km²) | % of gained AOH | % of lost AOH | name | Time period |
|---|---|---|---|---|---|---|
| 2022.5 | 2022.5 | 0 | 1 | 0 | *Bombus_inexspectatus* | T2 |
| 430.5 | 430.5 | 0 | 1 | 0 | *Bombus_konradini* | T2 |
| 225.4 | 225.4 | 0 | 1 | 0 | *Bombus_lucorum_aritzoensis* | T2 |
| 129.4 | 129.4 | 0 | 1 | 0 | *Bombus_monticola_alpestris* | T2 |
| 2491.8 | 469.3 | 264.1 | 0.7 | 0 | *Bombus_inexspectatus* | T3 |
| 430.5 | 0 | 0 | 0 | 0 | *Bombus_konradini* | T3 |
| 225.4 | 0 | 0 | 0 | 0 | *Bombus_lucorum_aritzoensis* | T3 |
| 129.4 | 0 | 129.4 | 0 | 0 | *Bombus_monticola_alpestris* | T3 |
| 11.6 | 11.6 | 0 | 1 | 0 | *Bombus_monticola_mathildis* | T3 |
| 2296.5 | 68.8 | 1524 | 1 | 0 | *Bombus_inexspectatus* | T4 |
| 430.6 | 0.1 | 0 | 1 | 0 | *Bombus_konradini* | T4 |
| 225.4 | 0 | 0 | 0 | 0 | *Bombus_lucorum_aritzoensis* | T4 |
| 11.6 | 0 | 0 | 0 | 0 | *Bombus_monticola_mathildis* | T4 |

| | | | | | | |
|---|---|---|---|---|---|---|
| 1095.3 | 1095.3 | 0 | 1 | 0 | *Bombus_alpinus_helleri* | T4 |
| 0.1 | 0.1 | 0 | 1 | 0 | *Bombus_brodmannicus_delmasi* | T4 |
| 1106.2 | 10.8 | 0 | 0.3 | 0 | *Bombus_alpinus_helleri* | T5 |
| 0.1 | 0 | 0 | 0 | 0 | *Bombus_brodmannicus_delmasi* | T5 |
| 772.5 | 0 | 0 | 0 | 0 | *Bombus_inexspectatus* | T5 |
| 546.4 | 115.7 | 2.7 | 1 | 0 | *Bombus_konradini* | T5 |
| 225.4 | 0 | 0 | 0 | 0 | *Bombus_lucorum_aritzoensis* | T5 |
| 11.6 | 0 | 0 | 0 | 0 | *Bombus_monticola_mathildis* | T5 |
| 1106.2 | 0 | 0 | 0 | 0 | *Bombus_alpinus_helleri* | T5 |
| 127.3 | 127.1 | 0 | 1 | 0 | *Bombus_brodmannicus_delmasi* | T5 |
| 772.5 | 0 | 463.9 | 0 | 0 | *Bombus_inexspectatus* | T5 |
| 544 | 0.3 | 0 | 1 | 0 | *Bombus_konradini* | T5 |
| 225.4 | 0 | 0 | 0 | 0 | *Bombus_lucorum_aritzoensis* | T5 |
| 11.6 | 0 | 0 | 0 | 0 | *Bombus_monticola_mathildis* | T5 |
| 2776.9 | 1670.7 | 616.9 | 0.9 | 0 | *Bombus_alpinus_helleri* | T6 |
| 652.8 | 525.6 | 8.1 | 1 | 0 | *Bombus_brodmannicus_delmasi* | T6 |
| 308.6 | 0 | 308.6 | 0 | 0 | *Bombus_inexspectatus* | T6 |
| 599.6 | 55.6 | 25.1 | 1 | 0 | *Bombus_konradini* | T6 |
| 255.2 | 29.7 | 43 | 1 | 0 | *Bombus_lucorum_aritzoensis* | T6 |
| 11.6 | 0 | 0 | 0 | 0 | *Bombus_monticola_mathildis* | T6 |
